# Cell-based analysis of *CAD* variants identifies individuals likely to benefit from uridine therapy

**DOI:** 10.1101/2020.03.11.987651

**Authors:** Francisco del Caño-Ochoa, Bobby G. Ng, Malak Abedalthagafi, Mohammed Almannai, Ronald D. Cohn, Gregory Costain, Orly Elpeleg, Henry Houlden, Ehsan Ghayoor Karimiani, Pengfei Liu, M. Chiara Manzini, Reza Maroofian, Michael Muriello, Ali Al-Otaibi, Hema Patel, Edvardson Shimon, V. Reid Sutton, Mehran Beiraghi Toosi, Lynne A. Wolfe, Jill A. Rosenfeld, Hudson H. Freeze, Santiago Ramón-Maiques

## Abstract

**Purpose:** Pathogenic autosomal recessive variants in *CAD*, encoding the multienzymatic protein initiating pyrimidine *de novo* biosynthesis, cause a severe inborn metabolic disorder treatable with a dietary supplement of uridine. This condition is difficult to diagnose given the large size of *CAD* with over 1000 missense variants and the non-specific clinical presentation. We aimed to develop a reliable and discerning assay to assess the pathogenicity of *CAD* variants and to select affected individuals that might benefit from uridine therapy.

**Methods:** Using CRISPR/Cas9, we generated a human *CAD*-knockout cell line that requires uridine supplements for survival. Transient transfection of the knockout cells with recombinant *CAD* restores growth in absence of uridine. This system determines missense variants that inactivate CAD and do not rescue the growth phenotype.

**Results:** We identified 25 individuals with biallelic variants in *CAD* and a phenotype consistent with a CAD deficit. We used the *CAD*-knockout complementation assay to test a total of 34 variants, identifying 16 as deleterious for CAD activity. Combination of these pathogenic variants confirmed 11 subjects with a CAD deficit, for whom we describe the clinical phenotype.

**Conclusions:** We designed a cell-based assay to test the pathogenicity of *CAD* variants, identifying 11 CAD deficient individuals, who could benefit from uridine therapy.

## INTRODUCTION

*CAD* encodes a multienzymatic cytoplasmic protein harboring four functional domains, each catalyzing one of the initial reactions for *de novo* biosynthesis of pyrimidine nucleotides: glutamine amidotransferase (GLN), carbamoyl-phosphate synthetase (SYN), aspartate transcarbamoylase (ATC) and dihydroorotase (DHO)^1-3^ (Figure 1). This metabolic pathway is essential for nucleotide homeostasis, cell growth and proliferation^4^. Defects in dihydroorotate dehydrogenase (DHODH) or UMP synthetase (UMPS), the enzymes catalyzing the next steps in the pathway after CAD, associate with severe human disorders (Miller syndrome [OMIM 263750]^5^ and orotic aciduria [OMIM 258900]^6^). In 2015, we identified a single individual with early infantile epileptic encephalopathy and two variants in *CAD*, one, an in-frame deletion of an exon and the second, a missense variant (p.R2024Q) in a highly conserved residue^7^. Metabolic analysis of subject fibroblasts showed impaired CAD activity-dependent incorporation of ^3^H-labeled aspartate into nucleic acids and nucleotide sugars, precursors for glycoprotein synthesis. Uridine supplements corrected this CAD-associated congenital disorder of glycosylation (CDG) [OMIM 616457], suggesting a simple potential treatment. In two subsequent reports, five affected individuals from four unrelated families with similar symptoms showed likely pathogenic variants in *CAD*, but no functional studies were done^8,9^. However, uridine treatment of three suspected individuals showed striking improvement, with cessation of seizures and significant progression from minimally conscious state to communication and walking. Recently, uridine triacetate (Xuriden) was approved by the FDA to treat hereditary orotic aciduria^10^; presumably, it could be used to treat affected individuals with CAD deficiency.

**Figure 1.**
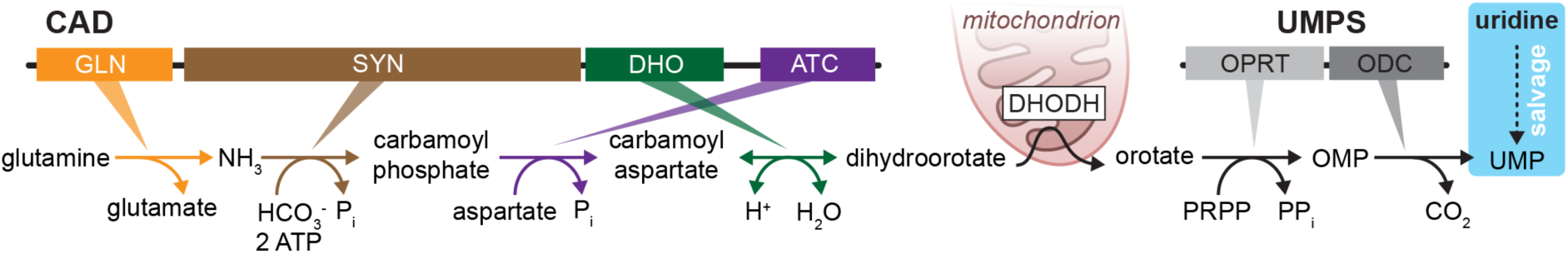
Schematic of the pathway for *de novo* biosynthesis of the pyrimidine nucleotide uridine 5-monophosphate (UMP). The initial enzymatic activities, glutaminase (GLN), carbamoyl phosphate synthetase (SYN), aspartate transcarbamoylase (ATC) and dihydroorotase (DHO) are fused into the multifunctional protein CAD. The next reaction after CAD is catalyzed by dihydroorotate dehydrogenase (DHODH), an enzyme anchored to the inner mitochondrial membrane. The last two steps are catalyzed by UMP synthase (UMPS), a bifunctional enzyme with orotate phosphoribosyl transferase (OPRT) and orotidine decarboxylase (ODC) activities. Alternatively, UMP can be obtained from uridine through salvage pathways (depicted in cyan).

The attractiveness of a simple therapy brought 25 suspected individuals to our attention for evaluation. Unfortunately, the metabolic labeling assay using ^3^H-labeled aspartate has a low resolution and a narrow dynamic range. To have a more reliable and discerning assay, we tested the ability of each variant to rescue growth of a human *CAD*-knockout cell line which requires uridine supplements for survival. Surprisingly, only 11 of 25 suspected individuals had pathologic variants and would potentially benefit from uridine supplements. We describe the development of this functional assay, the general clinical phenotype and analysis of these individuals. We caution about relying on current prediction programs to assess pathogenicity of variants for this large multifunctional enzyme.

## MATERIALS AND METHODS

### Clinical data

Informed consent was provided for all subjects in accordance with each clinician’s individual institution. For those individuals where samples were provided, written consent was provided for Sanford Burnham Prebys Medical Discovery Institute approved IRB-2014-038-17.

### CRISPR/Cas9 plasmid

pSpCas9 (BB)-2A-Puro (PX459) vector (Addgene), encoding Cas9, was digested with BbsI and purified with Qiaquick Gel Extraction kit (Qiagen). Complementary dsDNA oligonucleotides encoding sgRNA, designed to target the first exon of *CAD*, were purchased (Sigma) with 5’ overhangs complementary to the BbsI site and an extra G base to favor transcription^11^ (Table S1). The oligonucleotides were phosphorylated with T4 polynucleotide kinase (NEB), annealed and inserted in the linearized vector with T4 DNA ligase (NEB). The construct was amplified in TOP10 *E. coli* cells (ThermoFisher), verified by sequencing and purified with a Plasmid Midi kit (Qiagen).

### GFP-CAD plasmid

Enhanced green fluorescent protein (GFP) coding sequence was obtained by HindIII and KpnI digestion of pPEU2 vector (kindly provided by Dr. Nick Berrow, IRB Barcelona), and ligated into pCDNA3.1 (Promega) linearized with same restrictions enzymes. The resulting plasmid (pcDNA3.1-GFP) was verified by sequencing. Human *CAD* was PCR amplified from cDNA (Open Biosystems clone ID 5551082) using specific primers (Table S1) and ligated with In-Fusion (Clontech) into NotI linearized pcDNA3.1-GFP. The resulting plasmid (pcDNA3.1-GFPhuCAD) encodes an N-terminal histidine-tagged GFP followed in-frame by human CAD. Site-directed mutagenesis was carried out following the QuickChange protocol (Stratagene) and a pair of specific oligonucleotides (Table S1) and PfuUltra High-Fidelity DNA polymerase (Agilent).

### Generating a *CAD* knockout cell line

Human U2OS (bone osteosarcoma) cells were grown in DMEM (Lonza), 10% fetal bovine serum (FBS; Sigma), 2 mM L-glutamine (Lonza), and 50 U·ml^-1^ penicillin and 50 μg·ml^-1^ streptomycin (Invitrogen), at 5% CO_2_ and 37 °C. One day before transfection, 1.5–2 x 10^5^ U2OS cells in a final volume of 500 µl of medium were transferred to 24-well plates to reach approximately 50-80% confluence. For transfection, 2 µg of DNA in 50 µl of DMEM and 50 µl of FuGene6 transfection reagent (Promega) at 1mg·ml^-1^ in DMEM were incubated separately for 5 min at room temperature, and then mixed together and incubated at room temperature for an additional 10 min. The 100 µl mix was added to the wells drop by drop, followed by a 16 h incubation at 37°C and 5% CO_2_. 24 h post transfection, puromycin was added for one week to select transfected cells and enhance Cas9 cleavage. Media was supplemented with 30 µM uridine (Sigma) to allow growth of CAD deficient cells. Individual cells were isolated by serial dilution in 96-well plates, seeded into 24-well plates and expanded for 2-3 weeks. To identify CAD-deficient clones, a replica of the plate was grown in media with 10% fetal bovine macro-serum (FBM) without uridine, instead of FBS. FBM was prepared as reported^12^. In brief, 50 ml of heat-inactivated FBS were dialyzed against 1 L of tap water for 1 day at 4°C using SpectraPor #3 dialysis tubing with a molecular weight cutoff of 3,500 Da (Spectrum Laboratories, Inc., USA), supplemented with NaCl (9 g per liter), sterilized with a 0.22 µm filter and stored at −20 °C. Disruption of *CAD* was confirmed by Sanger sequencing. For this, exon 1 of *CAD* was PCR amplified with specific primers (Table S1), inserted in ZeroBlunt vector (Invitrogen) and sequenced with M13 primer. CAD-deficient cells were confirmed by Western blot and immunofluorescence microscopy using a monoclonal antibody (Cell Signaling Technology, #93925).

### Growth complementation assay

U2OS *CAD*-KO cells were transfected with wild-type (WT) or mutated pcDNA3.1-GFPhuCAD using FuGene6 as detailed above. One day after transfection, 1 × 10^5^ cells were seeded by duplicate in 24-well plates using media supplemented with 10% FBM (without uridine). Every 24 h, cells from one well were trypsinized and counted using a Countess II FL Automated Cell Counter (Thermo) or a Neubauer chamber. Doubling time was calculated using an online tool (http://www.doubling-time.com/compute.php).

## RESULTS

### Validation of a growth complementation assay in *CAD* knockout cells

We wanted to create a *CAD* knockout (KO) cell line that could be used to assess the pathogenicity of *CAD* variants. Using CRISPR/Cas9 technology, we knocked out *CAD* in human U2OS cells by selecting an isogenic clone that introduced a homozygous c.70delG frameshift [p.Ala24Profs*27] within exon 1 (Figure 2a-c). We verified by Western blot and immunofluorescence that *CAD*-KO cells do not express CAD (Figure 2c,d). As expected, these cells are unable to grow in absence of uridine, but proliferate at similar rate as WT cells in media supplemented with 30 µM exogenous uridine (Figure 2e). Next, we transiently transfected KO cells with a plasmid encoding human CAD fused to the enhanced green fluorescent protein (GFP) at the N-terminus (Figure 2f). *CAD*-KO cells expressing GFP-CAD proliferated in uridine-deprived conditions at a normal rate (doubling time ∼1 day), whereas cells transfected with GFP alone did not grow (Figure 2g).

**Figure 2.**
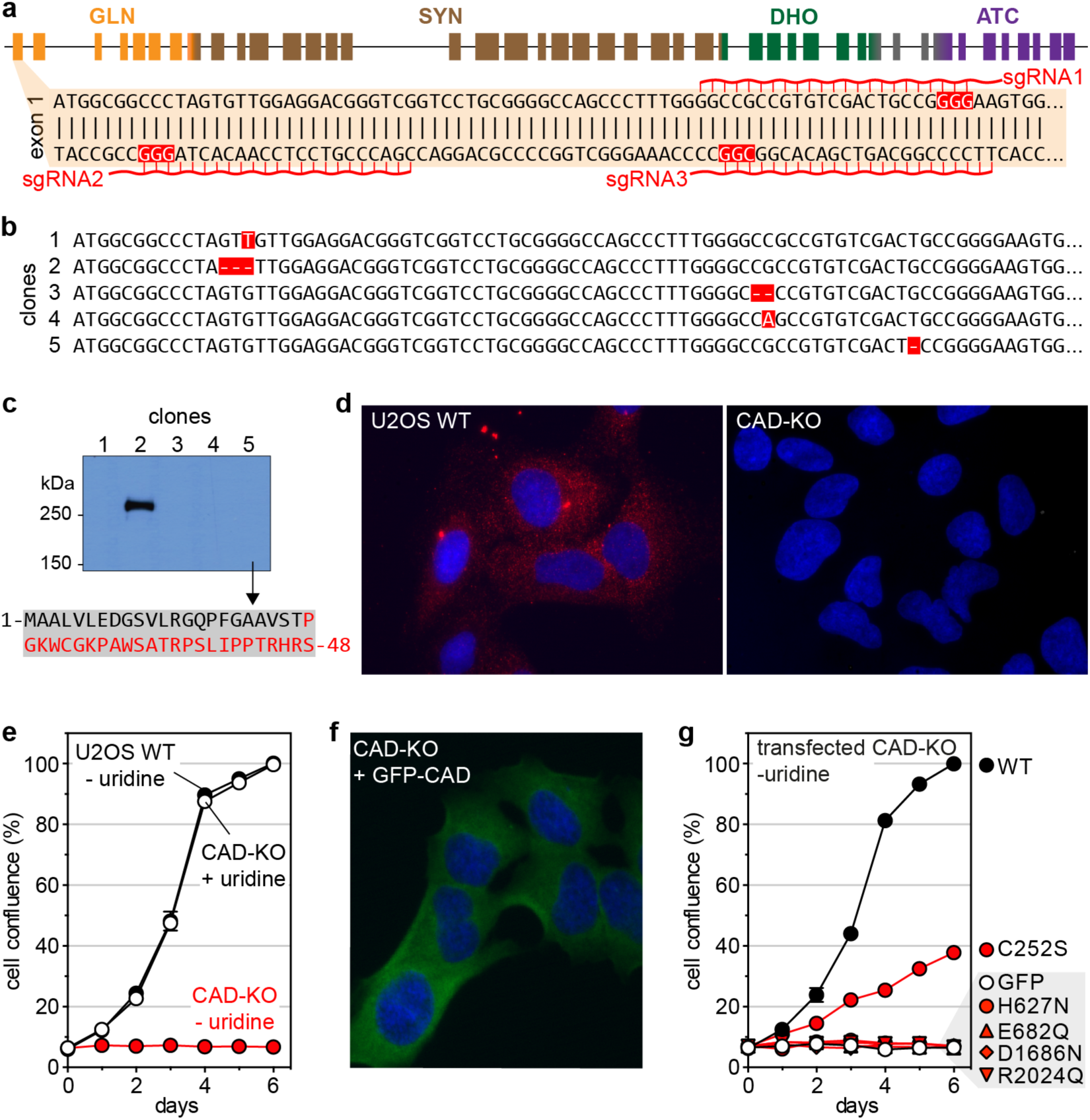
Using CRISPR/Cas9 to knockout *CAD* in U2OS cells. (**a**) Schematic representation of *CAD* locus, with 44 exons colored according to their respective functional domains; detail of the 5’ region of exon 1, indicating the single guide RNA (sgRNAs) with protospacer adjacent motif (PAM) sequences in red boxes. (**b**) Sequencing of five clones selected after CRISPR-Cas9 editing shows insertions and deletions (highlighted in red) in exon 1. (**c**) Expression of CAD in total lysates of clones shown in (**a**) analyzed by Western blot with a monoclonal antibody. Clone #5, chosen as the *CAD*-KO cell for further studies, produces an early truncated CAD protein of 48 residues with an incorrect sequence colored in red. (**d**) Immunofluorescence of WT and *CAD*-KO U2OS cells, using a monoclonal antibody against CAD (red signal) and nuclear labelling with Hoechst (blue signal). (**e**) Proliferation assay of *CAD*-KO cells in media with or without uridine, compared to the growth of WT cells. (**f**) Imaging of *CAD*-KO cells transiently transfected with GFP-CAD, using GFP fluorescent signal (green) and Hoechst (blue). (**g**) Transfection of GFP-CAD rescues the growth phenotype of *CAD*-KO in uridine-deprived media. Cells transfected with GFP alone do not proliferate. Cells transfected with GFP-CAD variants bearing well-characterized inactivating mutations in the SYN, DHO or ATC domains fail to proliferate without uridine, whereas the inactivation of the GLN domain (mutation C252S) allows limited growth.

To confirm that all four enzymatic activities of CAD were needed for *de novo* pyrimidine synthesis and cell growth in absence of uridine, we measured the proliferation of *CAD*-KO cells transfected with GFP-CAD bearing well-known inactivating mutations for each activity (Figure 2g). The transfected inactivated variants in the SYN (p.H627N, p.E682Q)^13,14^, DHO (p.D1686N)^15^ and ATC (p.R2024Q)^7,16^ domains failed to rescue the growth of *CAD*-KO cells. In turn, the GLN inactive mutant (p.C252S)^17^ showed a partial rescue, with transfected cells doubling every ∼2.5 days, suggesting that free-ammonia can, to some extent, contribute to the synthesis of carbamoyl phosphate (Figure 1).

### Identification and impact of potential CAD variants

Since *CAD* encodes a large protein with 2,225 amino acids covering 44 exons (Figure 2a), it is not surprising that all previously reported (n=6) affected individuals were identified using Next-Generation Sequencing (NGS)^7-9^. Likewise, using NGS we identified 25 potential CAD deficient individuals based on the presence of biallelic variants and a clinical phenotype similar to previously reported individuals (Table 1). Ultimately, we tested 34 variants of uncertain significance (VUS) in our validated knockout assay.

**Table 1.** Summary of *CAD* variants.

| ID<sup>a</sup> | cDNA<sup>b</sup> | Amino acid | SIFT<br>category | SIFT<br>value<sup>c</sup> | PolyPhen2<br>category | PolyPhen2<br>value<sup>c</sup> | CADD<br>PHRED<sup>c</sup> | KO rescue<br>results | gnomAD<br>carriers/alleles |
| --- | --- | --- | --- | --- | --- | --- | --- | --- | --- |
| Baylor - 001* | c.2156+5G>A | NA | NA | NA | NA | NA | 10.62 | NA | 4/251138 |
|  | c.4667A>C | p.K1556T | tolerated | 0.38 | possibly_damaging | 0.631 | 24.1 | **Pathogenic** | 1/251278 |
| Baylor - 002 | c.5147C>T | p.T1716M | deleterious | 0 | probably_damaging | 0.982 | 25.9 | Benign | 17/281494 |
|  | c.5561G>A | p.R1854Q | tolerated | 0.25 | benign | 0.186 | 23.8 | Benign | 15/282772 |
| Baylor - 003 | c.2372A>C | p.D791A | deleterious | 0.04 | possibly_damaging | 0.473 | 26.8 | Benign | NA |
|  | c.4487G>C | p.G1496A | deleterious | 0 | probably_damaging | 0.999 | 27 | Benign | NA |
| Baylor - 004 | c.713G>A | p.R238H | deleterious | 0.05 | benign | 0.104 | 18.27 | Benign | 109/282876 |
|  | c.1159G>A | p.G387S | tolerated | 0.67 | benign | 0 | 7.152 | Benign | 11/251452 |
| Baylor - 005 | c.4501T>A | p.C1501S | tolerated | 0.44 | possibly_damaging | 0.636 | 23.8 | Benign | 1/251356 |
|  | c.6556C>T | p.P2186S | deleterious | 0 | probably_damaging | 0.999 | 32 | **Pathogenic** | NA |
| Baylor - 006 | c.419A>G | p.Q140R | tolerated | 0.42 | benign | 0.029 | 18.14 | Benign | 2/282842 |
|  | c.5570G>A | p.R1857Q | tolerated | 0.17 | benign | 0.022 | 23.7 | Benign | 8/282788 |
| Baylor - 007 | c.943G>A | p.A315T | deleterious | 0.01 | probably_damaging | 0.971 | 26.4 | Benign | 4/251364 |
|  | c.5353C>T | p.R1785C | deleterious | 0 | probably_damaging | 0.994 | 28.5 | **Pathogenic** | 3/251030 |
| Baylor - 008 | c.785T>C | p.I262T | tolerated | 0.15 | benign | 0.185 | 22.7 | Benign | 7/282792 |
|  | c.3868G>A | p.G1290S | tolerated | 0.07 | benign | 0.058 | 16.78 | Benign | 31/282488 |
| Baylor - 009 | c.5147C>T | p.T1716M | deleterious | 0 | probably_damaging | 0.982 | 25.9 | Benign | 17/281494 |
|  | c.5561G>A | p.R1854Q | tolerated | 0.25 | benign | 0.186 | 23.8 | Benign | 15/282772 |
| Baylor - 010 | c.3649G>A | p.V1217I | deleterious | 0.02 | possibly_damaging | 0.483 | 25 | Benign | 3/251356 |
|  | c.4568C>T | p.A1523V | tolerated | 0.05 | benign | 0.391 | 23.6 | Benign | NA |
| Baylor - 011 | c.959A>G | p.N320S | deleterious | 0.02 | benign | 0.077 | 22.4 | **Pathogenic** | 9/282728 |
|  | c.2984C>G | p.S995C | deleterious | 0 | probably_damaging | 0.995 | 31 | Benign | NA |
| CDG - 0017 | c.1576G>A | p.G526R | deleterious | 0 | possibly_damaging | 0.657 | 26.7 | **Pathogenic** | 4/251224 |
|  | c.1576G>A | p.G526R | deleterious | 0 | possibly_damaging | 0.657 | 26.7 | **Pathogenic** | 4/251224 |
| CDG - 0104 | c.5959C>G | p.L1987V | deleterious | 0 | probably_damaging | 0.992 | 26.4 | **Pathogenic** | NA |
|  | c.5959C>G | p.L1987V | deleterious | 0 | probably_damaging | 0.992 | 26.4 | **Pathogenic** | NA |
| CDG - 0105 | c.5959C>G | p.L1987V | deleterious | 0 | probably_damaging | 0.992 | 26.4 | **Pathogenic** | NA |
|  | c.5959C>G | p.L1987V | deleterious | 0 | probably_damaging | 0.992 | 26.4 | **Pathogenic** | NA |
| CDG - 0111 | c.6329G>T | p.R2110L | tolerated | 0.2 | Benign | 0.046 | 16.83 | **Pathogenic** | 1/251314 |
|  | c.6329G>T | p.R2110L | tolerated | 0.2 | Benign | 0.046 | 16.83 | **Pathogenic** | 1/251314 |
| CDG - 0112 | c.3098G>A | p.R1033Q | deleterious | 0.01 | possibly_damaging | 0.537 | 31 | **Pathogenic** | 7/251346 |
|  | c.3098G>A | p.R1033Q | deleterious | 0.01 | possibly_damaging | 0.537 | 31 | **Pathogenic** | 7/251346 |
| CDG - 0117 | c.5957G>A | p.R1986Q | deleterious | 0.01 | probably_damaging | 0.992 | 33 | **Pathogenic** | 3/249932 |
|  | c.5957G>A | p.R1986Q | deleterious | 0.01 | probably_damaging | 0.992 | 33 | **Pathogenic** | 3/249932 |
| CDG - 0118 | c.6382G>A | p.E2128K | tolerated | 0.15 | possibly_damaging | 0.578 | 26.2 | **Pathogenic** | NA |
|  | c.6382G>A | p.E2128K | tolerated | 0.15 | possibly_damaging | 0.578 | 26.2 | **Pathogenic** | NA |
| CDG - 0122 | c.3512C>A | p.P1171Q | deleterious | 0 | probably_damaging | 0.936 | 28.4 | **Pathogenic** | 1/251476 |
|  | c.4315-1G>A | NA | NA | NA | NA | NA | 34 | NA | 1/31408 |
| CDG - 0123 | c.2995G>A | p.V999M | deleterious | 0 | probably_damaging | 1 | 29.8 | **Pathogenic** | NA |
|  | c.2995G>A | p.V999M | deleterious | 0 | probably_damaging | 1 | 29.8 | **Pathogenic** | NA |
| CDG - 0278 | c.98T>G | p.M33R | deleterious | 0 | benign | 0.223 | 24.9 | **Pathogenic** | 1/243848 |
|  | c.98T>G | p.M33R | deleterious | 0 | benign | 0.223 | 24.9 | **Pathogenic** | 1/243848 |
| CDG - 0443 | c.713G>A<br>Uniparental disomy Chr. 2 | p.R238H | deleterious | 0.05 | benign | 0.104 | 18.27 | Benign | 109/282876 |
| CDG - 1000 | c.4669C>G | p.L1557V | tolerated | 0.13 | benign | 0.003 | 21.9 | Benign | 62/282658 |
|  | c.6320C>G | p.P2107R | tolerated | 0.15 | benign | 0 | 16.49 | Benign | 2/282698 |
| CDG - 1001 | c.2386C>A | p.P796T | tolerated | 0.53 | benign | 0.039 | 21.1 | **Pathogenic** | 10/282430 |
|  | c.4735G>A | p.E1579K | tolerated | 0.26 | possibly_damaging | 0.624 | 23.7 | Benign | 5/250930 |
| CDG - 1046 | c.887G>A | p.G296E | deleterious | 0 | probably_damaging | 1 | 26.4 | **Pathogenic** | 6/251446 |
|  | c.2225G>A | p.R742Q | deleterious | 0.03 | benign | 0.414 | 25.3 | **Pathogenic** | NA |

<sup>a</sup> CAD-deficient subjects are denoted with gray background
<sup>b</sup> cDNA (NM\_004341.5), Uniprot ID (P27708)
<sup>c</sup> SIFT Value (Closer to 0 is damaging), Polyphen (Closer to 1 is damaging), CADD (20 puts variant in top 1% of deleterious variants, 30 in top 0.1%)
\*This individual was found to have both variants in *cis*.

To assess the damaging potential of variants found in subjects, we transfected *CAD*-KO cells with GFP-CAD bearing the clinical variants and monitored proliferation in uridine-deprived conditions (Figure 3a–d). Each newly constructed plasmid carrying an individual-specific variant required complete sequencing of the ∼8 kb *GFP-CAD* cDNA to ensure no additional changes were introduced during PCR. We also verified the efficiency of the transfection (>95%) and that the mutated proteins were being expressed by imaging the GFP fluorescence signal in the *CAD*-KO cells two days after transfection (data not shown).

**Figure 3.**
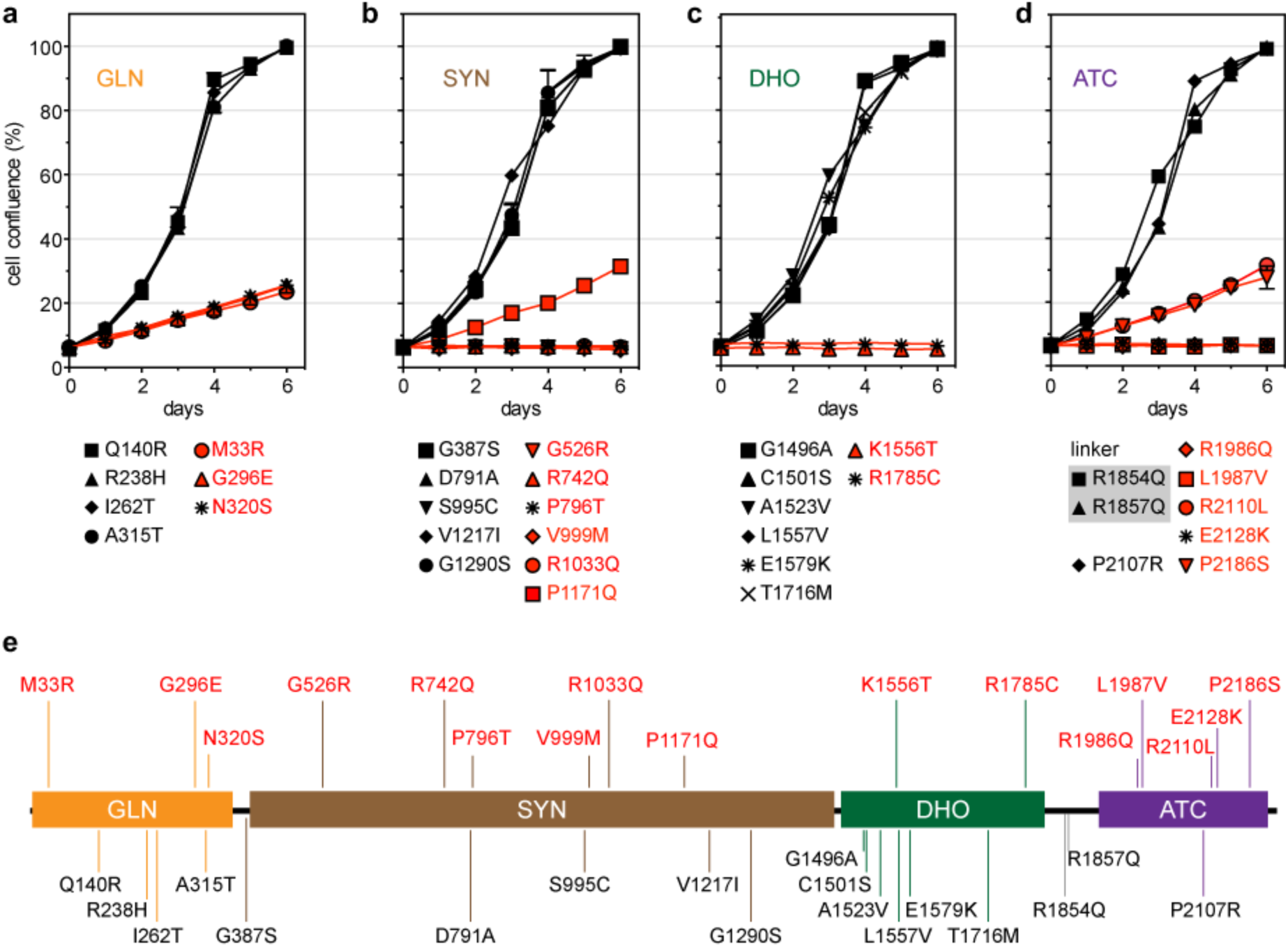
Assessing the pathogenicity of *CAD* variants. (**a-d**) Growth complementation assay of *CAD*-KO cells grown in absence of uridine and transfected with GFP-CAD bearing point mutations in the GLN (**a**), SYN (**b**), DHO (**c**) or ATC (**d**) domains. Mutations in the loop connecting the DHO and ATC domains are included in (**d**). Cell proliferation is represented as % confluence with respect to cells transfected with GFP-CAD WT. Each point represents the mean and standard deviation of three measurements, and all mutants were tested in at least two independent experiments. Mutations compromising CAD activity are colored in red. (**e**) Linear representation of CAD, mapping the inactivating (in red) and benign (in black) variants.

Three out of the seven variants found in the GLN domain, p.M33R, p.G296E and p.N320S, showed a partial rescue (Figure 3a). The doubling time was similar to the cells transfected with the GLN inactivating variant p.C252S (Figure 2g), indicating that these variants impair the GLN domain. On the other hand, cells transfected with SYN variants p.G526R, p.R742Q, p.P796T, p.V999M, and p.R1033Q failed to proliferate, whereas the variant p.P1171Q showed a partial rescue (Figure 3b). Out of the eight variants of the DHO domain tested, only two, p.K1556T and p.R1785C, failed to restore cell growth (Figure 3c). For the ATC variants, four mutations, p.R1986Q, p.L1987V and p.P2186S, failed to rescue the cells, whereas the p.E2128K allowed a partial rescue (Figure 3d). Finally, transfection with the two variants found at the linker between the DHO and ATC domains (p.R1854Q and p.R1857Q) restored normal growth (Figure 3d).

Based on these results, we concluded that the failure to rescue the growth phenotype of *CAD*-KO cells in absence of uridine indicates that 16 out of the 34 variants tested have a deleterious effect on CAD activity and therefore are pathogenic.

Interestingly, significant differences were seen when comparing the results of the KO assay to three popular *in silico* prediction programs (SIFT^18^, Polyphen2^19^, CADD^20^) (Table 1). All three prediction programs agreed with each other for 20/34 variants (59 % - 15/34 pathogenic, 5/34 benign variants). Yet only 38 % (13/34) (9/34 pathogenic, 4/34 benign) of the variants agreed in all three prediction programs and the complementation assay. We used a CADD PHRED score of above 20, which places a variant in the top 1% deleterious variants in the human genome, as potentially pathogenic. Below 20 we considered likely benign.

The mechanisms of inactivation of the pathogenic variants will be described in a separate study.

### Clinical

To date, only six affected individuals from five unrelated families have been identified with CAD deficiencies^7-9^. The clinical presentation of these individuals is general in nature, but all showed varying severity of neurological involvement including developmental delays and/or seizures. Furthermore, all were reported to have hematological abnormalities including abnormal red blood cells (anisopoikilocytosis) and anemia. Two of the six are reported to be deceased, while the remaining four were placed on uridine supplementation.

In this study, we identified 25 individuals with biallelic variants in *CAD*, who presented with a phenotype potentially consistent with CAD deficiency. We used the *CAD*-KO complementation assay described above to determine the pathogenicity of each variant identified and ultimately confirmed eleven CAD-deficient subjects (Table 1, Figure 3e).

Detailed clinical information was available and provided for ten of the eleven confirmed individuals (Figure 4). Consistent with the initial CAD deficient individuals^7,8^, all ten individuals presented here showed varying neurological abnormalities. All (10/10, 100 %) had intellectual and development delays, while 9/10 (90 %) had seizure activity. Gastrointestinal complications ranging from feeding problems, reflux and recurrent vomiting were seen in half (5/10) of the individuals. Facial dysmorphism, hypotonia and ataxia were also seen in half of those affected. While the sample size of previously identified subjects is small, 5/5 (100 %) did show hematological abnormalities. In contrast, our cohort reported only 4/10 (40 %) with these. Less affected systems included the skeletal (3/10) and cardiac (2/10).

**Figure 4.**
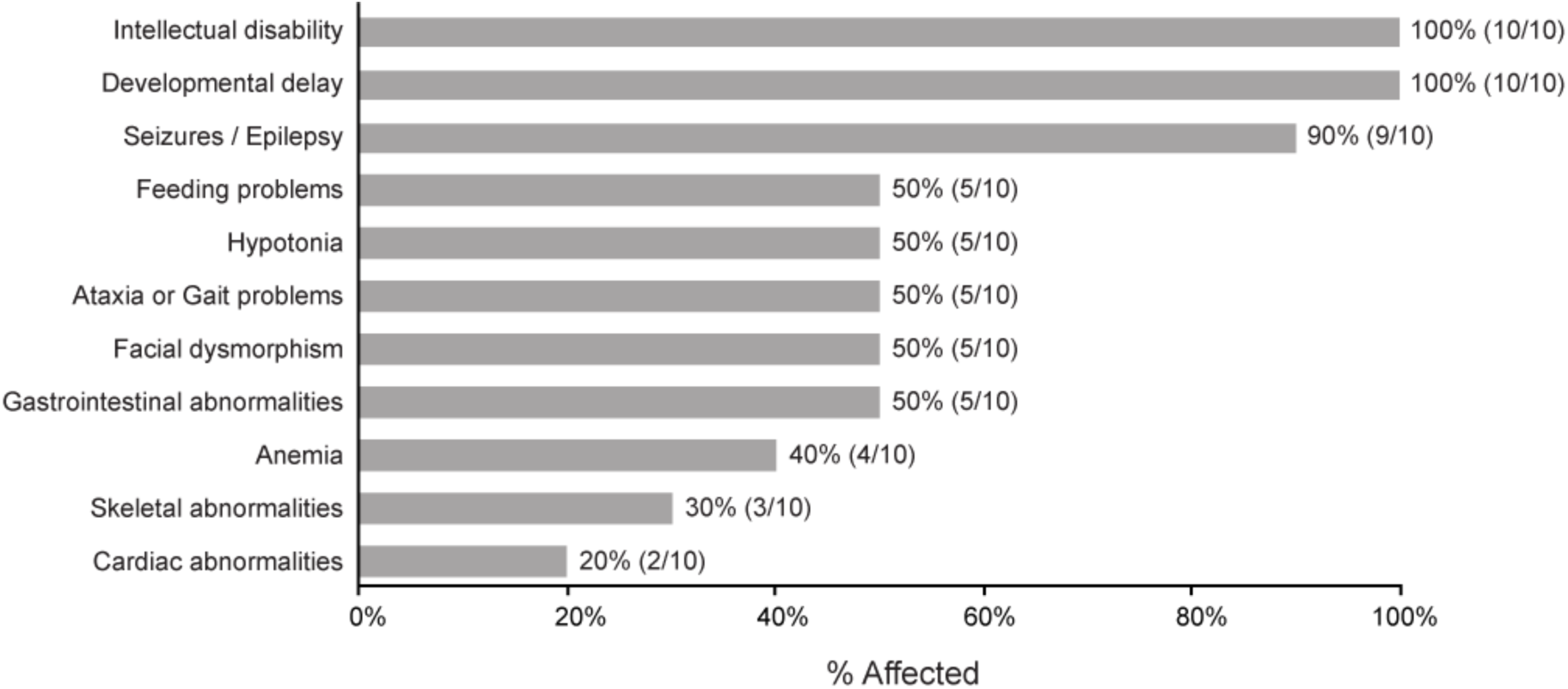
Clinical summary for ten unreported CAD-deficient individuals. Clinical information for 10 of the available subjects was collected and summarized as % affected.

In our cohort, one individual was noted to have passed away (CDG-0118). However, four families (0017, 0104, 0118, 0123) were noted to have a family history of multiple affected siblings with a similar presentation. From these four families, three had at least one sibling with a similar disorder who expired.

Due to the lack of detailed clinical information, CDG-0117 was not included in the final summary. However, he was noted to have structural brain abnormalities and a family history significant for premature death in two affected female siblings. Importantly, genomic DNA was available for one of the two deceased siblings and was found to also carry the same homozygous c.5957G>A [p.R1986Q] *CAD* variant.

One family (CDG-0112) had a dual diagnosis of CAD deficiency and a recessive intellectual developmental disorder with cardiac arrhythmia (OMIM 617173). Within this family, both affected siblings harbored a homozygous pathogenic c.249+3A>G [p.Asp84Valfs31*] variant in *GNB5*^21^, but only the male sibling carried the pathogenic homozygous c.3098G>A [p.R1033Q] variant in *CAD*. Given the clinical similarities of these two disorders, especially the neurological features, we cannot determine which symptoms are due to specifically the CAD variant alone.

## DISCUSSION

The prospect of a simple, non-toxic therapy for a potentially lethal disorder excites all stakeholders: patients, caretakers, physicians and scientists. Identifying the first CAD-deficient individual and showing that uridine corrects cellular defects set the stage for the highly successful use of uridine in two CAD-deficient individuals^7,8^. As a result, and given the non-specific clinical presentation of CAD-deficient individuals, we received many requests to test subject fibroblasts in a functional assay that involves labelling cells with ^3^H-aspartate to measure the CAD-dependent contribution to *de novo* pyrimidine synthesis (Figure 1). However, the assay has a limited dynamic range (∼2 fold) and many determinations left us ambivalent and uncertain about the diagnosis. Thus, a new robust and reliable biochemical assay was required to evaluate the pathogenicity of *CAD* variants.

We designed a *CAD* knockout cell line whose growth was dependent on added uridine (Figure 2) and then tested each variant for its ability to rescue uridine-independent growth (Figure 3a-d). Most of the variants either fully rescued growth, meaning the variants were benign, or were unable to rescue growth completely, showing they were pathologic variants. Only a few showed partial rescue, which we interpret to mean a damaging variant that decreases, but does not eliminate the activity. When each variant was combined based on individual-specific genotyping, we determined which individuals indeed had a CAD deficiency and therefore predict which ones would benefit from uridine therapy (Figure 3e and Table 1). This is a stringent prediction based on each single variant. It does not test the specific combination of alleles found in each individual, but we assume the combination of two variants would not cancel each other to generate a fully capable CAD protein. If this were the case, it is unlikely that the individuals themselves would show the expected clinical phenotype. Surprisingly only 11 of the 25 suspected individuals appear to be authentic cases based on this functional assay.

We also compared our assay results to three prediction programs designed to assess the pathogenicity of each variant (Table 1). There was considerable disagreement between the programs for many variants, and the programs produced both false positive and false negative results. Based on these findings, we suggest that any suspected CAD cases first be validated using this (or similar) biochemical assay. And it is likely that more putative CAD deficient cases will be suspected, since *CAD* has ∼1,020 missense rare variants in the public gnomAD browser (Ver2.1.1) database^22^ (accessed 2020.1.23 with 125,748 exomes and 15,708 genomes). Some families may choose to start uridine therapy without benefit of these results. That is certainly possible since the uridine is available to families and subjects over the internet. Barring the consumption of impure products, uridine is unlikely to be harmful. On the other hand, using uridine supplements in unconfirmed subjects may offer false hopes and complicate the interpretation of successful uridine therapy.

## ACKNOWLEDGEMENTS

This work was supported by The Rocket Fund, by R01DK99551 to HHF, and by MICIU grant BFU2016-80570-R and RTI2018-098084-B-100 (AEI/FEDER, UE). The University of Washington Center for Mendelian Genomics for exome sequencing and analysis of CDG - 0117. We would like to thank all the families for providing biological samples and their continued support. We would like to also thank the clinicians who provided information for individuals that were determined not to be CAD deficient.

## Supplementary Information

**Table S1.**
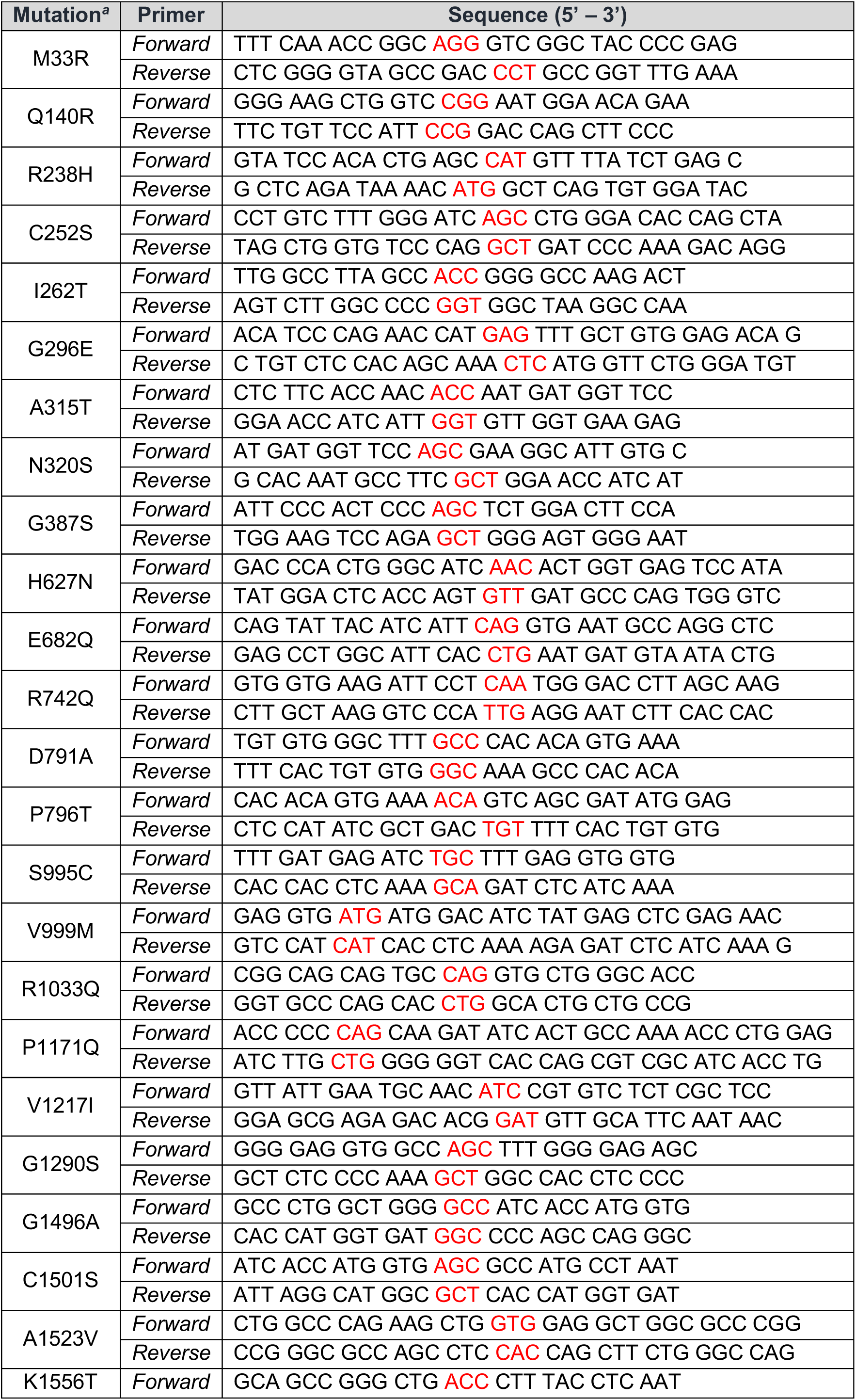
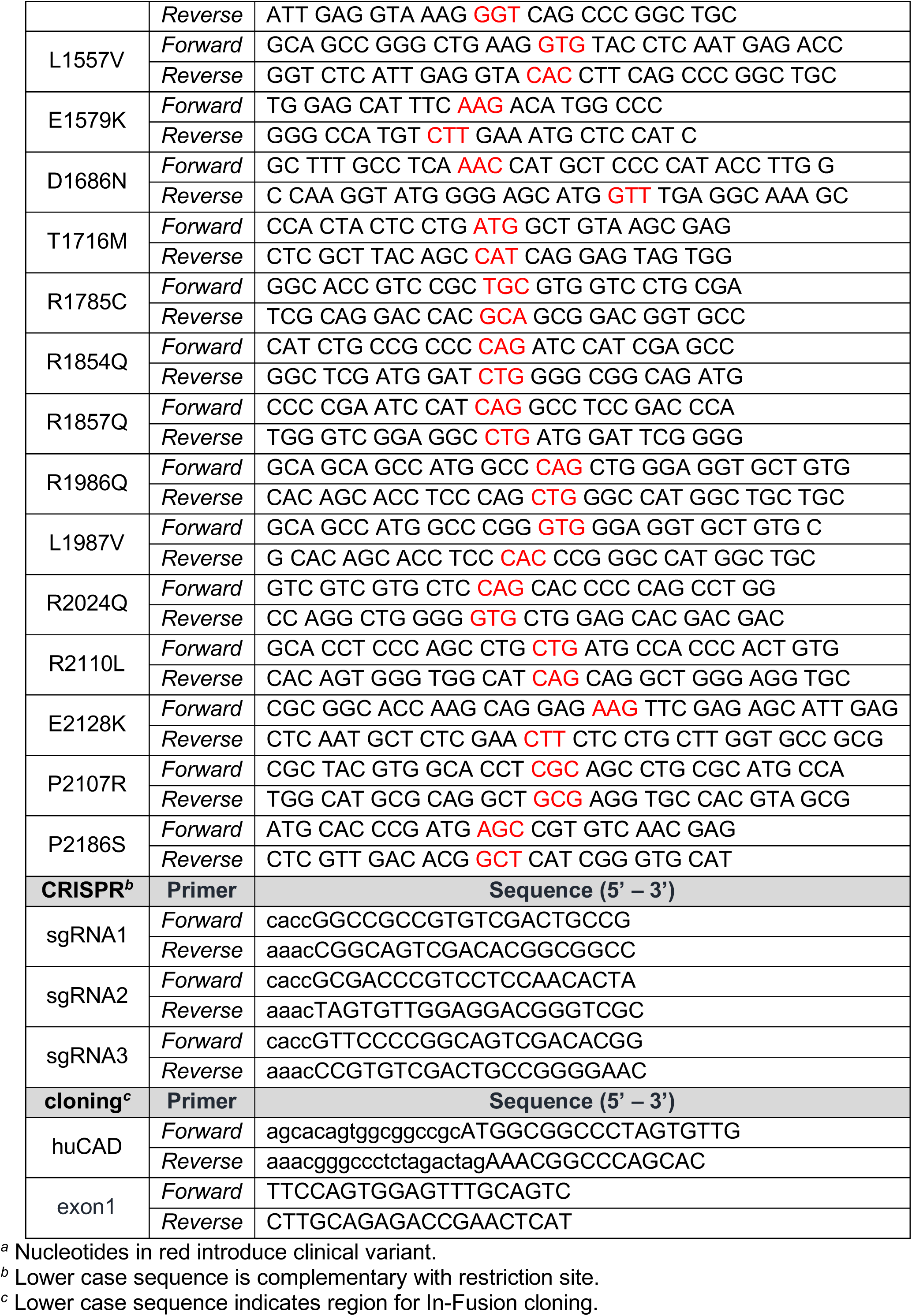
Oligonucleotides used for site-directed mutagenesis, CRISPR/Cas9 editing and cloning.

